# Endogenous neuronal DNA double-strand breaks are not sufficient to drive brain aging and neurodegeneration

**DOI:** 10.1101/2024.10.22.619740

**Authors:** Sarah Cohen, Laura Cheradame, Karishma J. B. Pratt, Sarah Collins, Ashlie Barillas, Annika Carlson, Vijay Ramani, Gaëlle Legube, Saul A. Villeda, R. Dyche Mullins, Bjoern Schwer

## Abstract

Loss of genomic information due to the accumulation of somatic DNA damage has been implicated in aging and neurodegeneration^1–3^. Somatic mutations in human neurons increase with age^4^, but it is unclear whether this is a cause or a consequence of brain aging. Here, we clarify the role of endogenous, neuronal DNA double-strand breaks (DSBs) in brain aging and neurodegeneration by generating mice with post-developmental inactivation of the classical non-homologous end-joining (C-NHEJ) core factor Xrcc4 in forebrain neurons. Xrcc4 is critical for the ligation step of C-NHEJ and has no known function outside of DSB repair^5,6^. We find that, unlike their wild-type counterparts, C-NHEJ-deficient neurons accumulate high levels of DSB foci with age, indicating that neurons undergo frequent DSBs that are typically efficiently repaired by C-NHEJ across their lifespan. Genome-wide mapping reveals that endogenous neuronal DSBs preferentially occur in promoter regions and other genic features. Analysis of 3-D genome organization shows intra-chromosomal clustering and loop extrusion of neuronal DSB regions. Strikingly, however, DSB accumulation caused by loss of C-NHEJ induces only minor epigenetic alterations and does not significantly affect gene expression, 3-D genome organization, or mutational outcomes at neuronal DSBs. Despite extensive aging-associated accumulation of neuronal DSBs, mice with neuronal Xrcc4 inactivation do not show neurodegeneration, neuroinflammation, reduced lifespan, or impaired memory and learning behavior. We conclude that the formation of spontaneous neuronal DSBs caused by normal cellular processes is insufficient to cause brain aging and neurodegeneration, even in the absence of C-NHEJ, the principal neuronal DSB repair pathway.

## MAIN

The brain has been proposed to be especially vulnerable to DNA damage and genomic alterations due to its high metabolic requirements, transcriptional activity, the long lifespan of neurons, and its limited regenerative capacity^7^. Neuronal activity threatens genome integrity as it can induce DSBs in genes critical for neuronal plasticity, learning, and memory^8–11^. Recent studies have proposed that excessive DSB formation may promote neurodegenerative disorders *via* the accumulation of somatic mutations, epigenetic landscape erosion, alterations in 3-D genome organization, and induction of neuroinflammation^12–14^.

These previous studies relied on artificial, ‘exogenous’ mechanisms of inducing DNA damage, including expression of the endonuclease *I-PpoI* or overexpression of p25^12–14^. Such non-physiological sources of DNA damage differ from normally occurring, ‘endogenous’ DNA damage in several key features, such as: the number of DSBs and the kinetics of their accumulation; the distribution of DSB sites within the genome; and the mechanism-specific molecular context. We, therefore, wondered whether conclusions from the previous studies are relevant to DNA damage that accumulates *via* endogenous, age-related mechanisms.

Postmitotic neurons are thought to primarily rely on the classical non-homologous end-joining (C-NHEJ) pathway for DSB repair^15^. To elucidate the impact of endogenous DNA double-strand breaks on neuronal function, aging, and neurodegeneration (**Fig. 1a**), we generated mice with postnatal neuron-specific inactivation of C-NHEJ by crossing a well-established conditional *Xrcc4* knockout allele^16^ to mice expressing *calcium/calmodulin-dependent protein kinase II alpha* (*Camk2a*) promoter-driven Cre recombinase^17^. This approach enables conditional *Xrcc4* knockout (*Xrcc4* nKO) in postmitotic excitatory neurons of the cortex and hippocampus during the third postnatal week^17^ and avoids confounding effects of neurodevelopmental *Xrcc4* ablation^18^.

**Figure 1.**
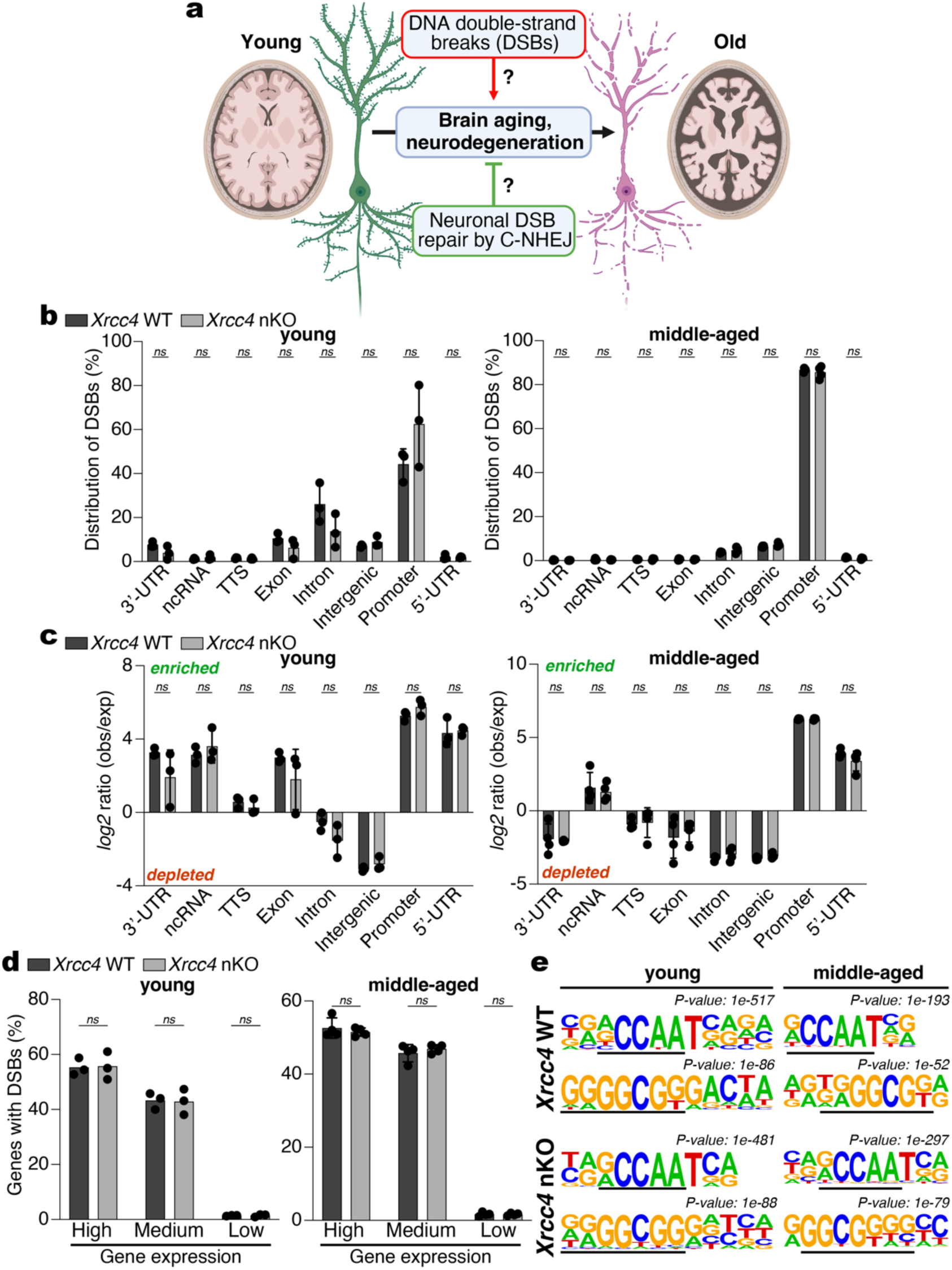
Endogenous DSBs are enriched in transcribed genes in the cerebral cortex of wild-type and neuron-specific *Xrcc4* knockout mice. **a,** Schematic of the potential interplay between endogenous neuronal DSBs, C-NHEJ, brain aging, and neurodegeneration. **b**, Genomic distribution of sBLISS consensus peaks in young (*n* = 3 mice per genotype; 3 months) or middle-aged (*n* = 4 mice per genotype; 13.1 ± 1.1 months) wild-type (WT) and *Xrcc4* nKO mice. **c**, Distribution of sBLISS peaks across the indicated genome annotations. The *log2* enrichment ratio of the observed *versus* expected genomic distribution is shown. **d**, Expression levels of genes with DSBs determined by RNA-seq analysis of the cerebral cortex from young or middle-aged, wild-type or *Xrcc4* nKO mice, respectively. High, >1,000 transcripts per million reads (TPM); medium, 10–1,000 TPM; low, 1–10 TPM. Data in **b–d** represent mean ± s.d; *ns*, not significant; multiple unpaired *t*-tests with two-stage step-up (Benjamini, Krieger, and Yekutieli) multiple comparison and FDR = 1%). **e**, *De novo* motif analysis of sBLISS consensus peaks identified in young or middle-aged, wild-type or *Xrcc4* nKO cerebral cortex, respectively.

Xrcc4 functions exclusively in the final ligation step of DSB repair *via* C-NHEJ^6^. We thus hypothesized that *Xrcc4* ablation would not alter the genomic location of spontaneous, endogenous DSBs in neurons. To test this, we mapped the genome-wide distribution of spontaneously occurring, endogenous DSBs by performing *in-suspension Breaks Labeling In Situ and Sequencing* (sBLISS) on isolated nuclei from the cortex and hippocampus of wild-type or *Xrcc4* nKO mice (Fig. 1b and **Extended Data Fig. 1**). This revealed that DSBs in young and middle-aged wild-type or *Xrcc4* nKO neurons localize preferentially to promoters and gene bodies and are depleted in introns and intergenic regions **(Fig. 1b,c)**. DSBs occurred predominantly in genes with high or medium expression levels in the cerebral cortex (**Fig. 1d)**, indicating that transcription-associated processes are a significant source of endogenous DSBs in the cerebral cortex.

To further assess determinants of neuronal DSBs, we analyzed the genomic sequences surrounding sBLISS peaks present in two of three biological sBLISS replicates from young mice, and three of four biological sBLISS replicates from middle-aged mice, respectively. This revealed a strong enrichment for a common promoter element (CCAAT)^19^ motif and a G-rich transcription factor binding motif (GGCGGG) in young and middle-aged mice of both genotypes (**Fig. 1e)**. GC-rich sequences can form G-quadruplex secondary structures and have been implicated in R-loop-mediated DSB formation in promoter regions^20^, suggesting that these features may enhance genomic fragility of these regions in neurons. These findings show that endogenous DSBs occur preferentially at promoters and 5’-untranslated regions (5’-UTRs) of robustly transcribed genes with specific DNA motifs. Consistent with the exclusive role of Xrcc4 in the ligation step of C-NHEJ-mediated DSB repair^5,6^, *Xrcc4* ablation did not affect the genomic distribution of DSBs. Thus, our experimental approach is ideal for revealing the neurobiological implications of neuronal DSB formation and repair and avoids issues associated with generating non-physiological DSBs^12^.

Next, we assessed if neuronal DSBs accumulate with age and at different rates in wild-type and *Xrcc4* nKO mice. We performed immunofluorescence microscopy of brain sections of wild-type and *Xrcc4* nKO mice against two established DSB markers, γH2AX and 53BP1^21^. Approximately 10% of neurons in the hippocampus (*Dentate gyrus*) and cerebral cortex contained, on average, one γH2AX focus in both young and old wild-type mice, suggesting that neurons undergo frequent DSBs but that these do not accumulate with age (**Fig. 2a,b** and **Extended Data Fig. 2a).** In stark contrast, cortical and hippocampal neurons lacking C-NHEJ showed increased DSB foci at a young age, which further increased with age (**Fig. 2a,b** and **Extended Data Fig. 2a**). Strikingly, up to 97% of aged C-NHEJ-deficient neurons contained DSB foci, with most of these containing three or more DSB foci per neuron (**Fig. 2a,b** and **Extended Data Fig. 2a).** We did not detect substantial numbers of DSB foci in non-neuronal brain cells at either age (**Extended Data Fig. 2b,c).** These findings show that neurons frequently undergo DSBs from endogenous sources across their lifespan, which are normally repaired efficiently by C-NHEJ.

**Figure 2.**
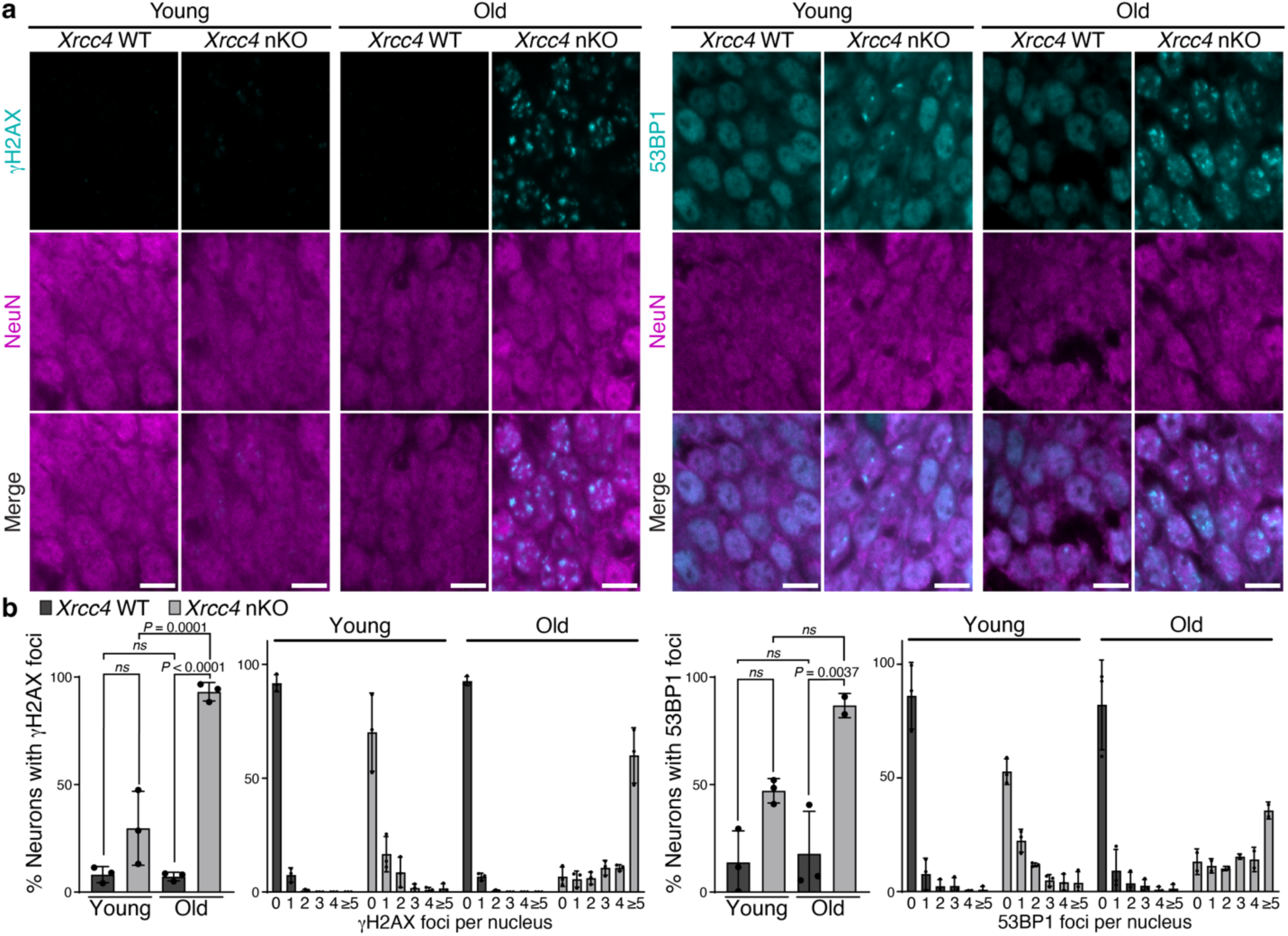
Neuronal C-NHEJ suppresses aging-associated DSB accumulation. **a**, Representative images of DSB response foci marked by γH2AX (*left*) and 53BP1 (*right*) in hippocampal (*Dentate gyrus)* sections of young (6.1 ± 1.3-month-old) and old (21.4 ± 2.5-month-old) wild-type or *Xrcc4* nKO mice; scale bars, 10 μm. **b**, Quantification of the fraction of NeuN^+^ hippocampal neurons (*Dentate gyrus)* with DSB response foci and distribution of the number of foci per neuron based on γH2AX (*left graphs*) or 53BP1 (*right graphs*) staining. Data represent mean ± s.d.; individual points show the mean from two fields per mouse. *P*-values were determined by one-way ANOVA and Tukey’s *post-hoc* multiple comparisons test; *ns*, not significant.

Recent studies relying on non-physiological, endonuclease-based DSBs have implicated DSBs as a cause of aging *via* the loss of epigenetic information^12^. To assess if endogenous neuronal DSBs may cause brain aging by eroding the epigenetic landscape, we assessed active (histone H3 lysine 4 tri-methylation (H3K4me3), histone H3 lysine 27 acetylation (H3K27ac)) and repressive (histone H3 lysine 27 tri-methylation (H3K27me3)) chromatin marks in the cerebral cortex and hippocampus of young wild-type or *Xrcc4* nKO mice by *Cleavage Under Targets & Release Using Nuclease* (CUT&RUN; **Fig. 3a–c**). We also measured chromatin accessibility *via Assay for Transposase-Accessible Chromatin by sequencing* (ATAC-seq; **Fig. 3a–c**). DSB sites identified by sBLISS were highly enriched in open chromatin with active histone marks in wild-type and *Xrcc4* nKO mice (**Fig 3a–c**). We noted a small but significant decrease of H3K4me3 and H3K27me3 and an increase in H3K27ac signal around DSB sites in *Xrcc4* nKO mice (**Fig. 3b,c**). However, chromatin accessibility around DSB sites was not significantly altered (**Fig. 3b,c**). Our results suggest that C-NHEJ deficiency leads to a small but significant alteration of the chromatin landscape around endogenous DSBs.

**Figure 3.**
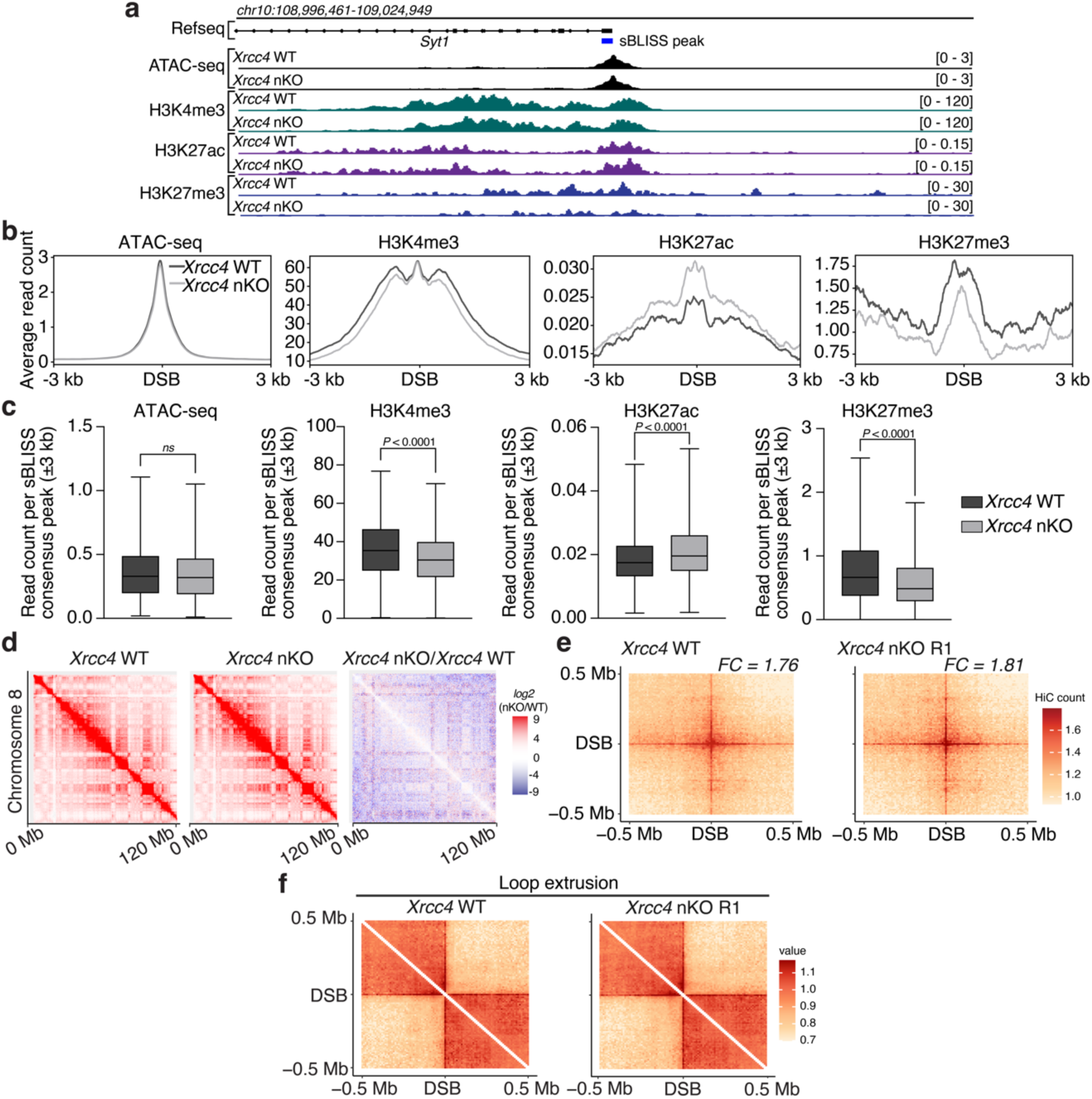
Neuronal C-NHEJ inactivation causes minor alterations to the epigenome but does not affect chromatin accessibility at DSB regions or alter 3-D genome organization. **a**, Genomic tracks showing chromatin accessibility (ATAC-seq), H3K4me3, H3K27ac, and H3K27me3 signal in the vicinity of an sBLISS consensus peak in *Syt1* (blue bar) in the cerebral cortex. The ATAC-seq signal shown is from four biological replicates per genotype (*n* = 4 mice; 5.2 ± 1.1 months). H3K4me3, H3K27ac, and H3K27me distribution were determined by CUT&RUN using *S. cerevisiae* spike-in DNA for normalization. The spike-in-normalized aggregate signal from biological replicates is shown (H3K4me3, *n* = 2 mice per genotype; H3K27ac, *n* = 3 mice per genotype; H3K27me3, *n* = 2 mice per genotype; 5.2 ± 1.1 months). **b**, Chromatin accessibility (ATAC-seq) and H3K4me3, H3K27ac or H3K27me3 CUT&RUN signal within 3 kb of the center of sBLISS consensus peaks (DSB) detected in wild-type and *Xrcc4* nKO cerebral cortex. **c**, Quantification of chromatin accessibility, H3K4me3, H3K27ac, or H3K27me3 signal within 3 kb of the center of sBLISS consensus peaks detected in wild-type and *Xrcc4* nKO cerebral cortex without outliers (identified with ROUT Q = 1%). Whiskers show minimum and maximum values; top and bottom edges of boxplots correspond to 25th and 75th percentiles, respectively; horizontal lines indicate the median. *P*-values were determined by the Kolmogorov-Smirnov test; *ns*, not significant. **d**, Hi-C contact matrices of chromosome 8 in wildtype (*left*) and *Xrcc4* nKO (biological replicate R1, *middle*) and differential Hi-C contact matrix (*log2*[*Xrcc4* nKO/WT], *right*) in cerebral cortex are shown; see **Extended Data Figure 3** for additional replicates. **e**, Mean aggregate peak analysis of intra-chromosomal DSB consensus peak interactions plotted on a 500-kb window (10-kb resolution) in wild-type and *Xrcc4* nKO cerebral cortex. *FC*, fold change calculated between the central pixel and a square of 3×3 pixels on the bottom left corner of the matrix. **f**, Aggregate Hi-C contact matrix plotted on a 500-kb window (10-kb resolution) centered on sBLISS consensus peaks in wild-type and *Xrcc4* nKO cerebral cortex.

Although a recent study suggested that neuronal DSBs cause 3-D genome disruptions in the context of neurodegeneration^13^, our Hi-C analysis of global 3-D genome organization did not reveal significant differences between the cerebral cortex of wild-type and *Xrcc4* nKO mice (**Fig. 3d** and **Extended Data Fig. 3a**). We did, however, detect intra-chromosomal clustering of genomic regions containing DSB sites (**Fig. 3e** and **Extended Data Fig. 3b**) consistent with observations made in dividing human cells^22^. Clustering of DSBs was present in the cerebral cortex of wild-type and *Xrcc4* nKO mice to a similar extent, indicating that it is not affected by the loss of neuronal C-NHEJ (**Fig. 3e** and **Extended Data Fig. 3b**). Notably, aggregate Hi-C maps of *cis* interaction frequencies centered on DSB sites showed a pattern associated with loop extrusion (**Fig. 3f** and **Extended Data Fig. 3c**)^23^. These findings reveal that endogenous neuronal DSBs cluster and undergo loop extrusion; however, we conclude that the accumulation of endogenous DSBs in the absence of C-NHEJ is not sufficient to disrupt neuronal 3-D genome organization.

A dramatic enhancement in the number of endogenous neuronal DSBs is not sufficient to cause substantial age-associated genomic alterations. We analyzed bulk RNA-seq data from the cerebral cortex of 16-month-old wild-type or *Xrcc4* nKO mice for chimeric reads indicative of structural variants. Levels of gene fusion transcripts were low in both wild-type and *Xrcc4* nKO cortex (**Fig. 4a**), suggesting that structural variants arising from DSBs, such as inter-and intra-chromosomal translocations, are not common in the cerebral cortex. To further examine the potential mutagenic outcomes of endogenous neuronal DSBs, we performed targeted amplicon sequencing^11,24^ of 12 regions surrounding DSBs detected in both wild-type and *Xrcc4* nKO neurons. Surprisingly, neurons from old wild-type or *Xrcc4* nKO mice showed only low levels of insertions and deletions (**Fig. 4b**).

**Figure 4.**
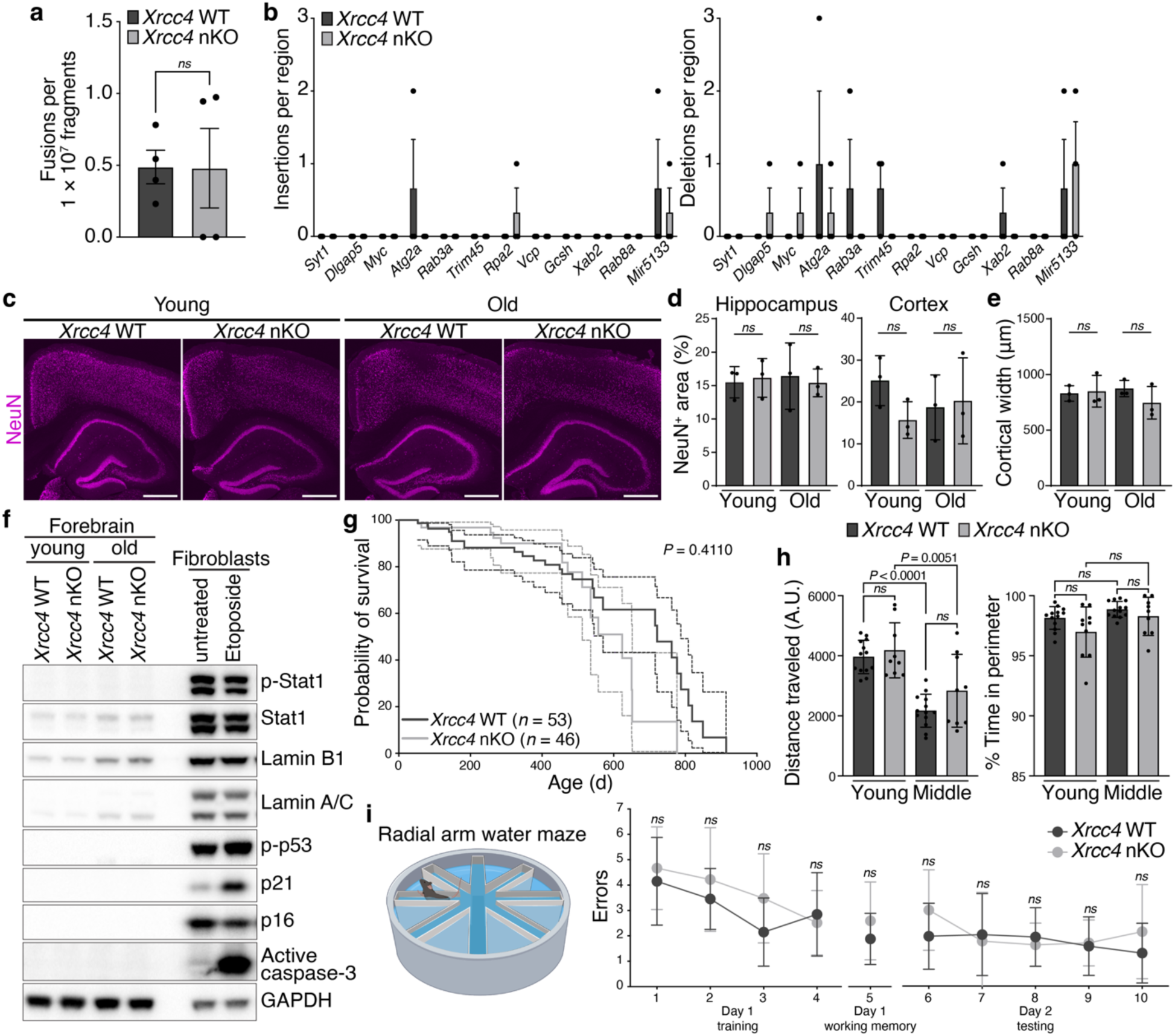
Neuronal DSBs caused by endogenous processes are not sufficient to cause brain aging or neurodegeneration, even when C-NHEJ is inactivated. **a**, Number of gene fusions detected in 15.9 ± 1.0 month-old wild-type and *Xrcc4* nKO mice (*n* = 4 mice per genotype). Data represent mean ± s.e.m; *ns*, not significant (unpaired, two-tailed *t*-test). **b**, Insertions (*left*) and deletions (*right*) determined by targeted amplicon sequencing of 12 distinct DSB regions in old (25.3 ± 5.2 months) wild-type or *Xrcc4* nKO mice (*n* = 3 mice per genotype). **c**, Representative images of NeuN-stained coronal brain sections (cortex and hippocampus) from young (6.1 ± 1.3 months) and old (21.4 ± 2.5 months) wild-type or *Xrcc4* nKO mice; scale bars, 500 μm. **d**, Quantification of NeuN-positive area in the hippocampus (*Dentate gyrus*, *Cornu ammonis* (CA)1, CA2, and CA3; *left*) and cortex (*right*) of young (6.1 ± 1.3 months) and old (21.4 ± 2.5 months) wild-type or *Xrcc4* nKO mice (*n* = 3 mice per genotype). **e**, Quantification of cortical width in young (6.1 ± 1.3 months) and old (21.4 ± 2.5 months) wild-type or *Xrcc4* nKO mice (*n* = 3 mice per genotype). Data in **d** and **e** represent mean ± s.d.; one-way ANOVA and Tukey’s *post-hoc* multiple comparison tests. **f**, Representative immunoblot analysis of proteins associated with inflammation (interferon-stimulated gene Stat1 expression and phosphorylation), senescence (p21, p16, lamin A/C, and lamin B1), apoptosis (active caspase-3), and checkpoint activation (phosphorylated p53) in young (3.2 months) and old (23.4 ± 0.8 months) wild-type or *Xrcc4* nKO forebrain (pooled cortex and hippocampus). Whole-cell extracts from SV40 large T antigen-transformed murine tail fibroblasts with or without etoposide treatment (10 μM for 24 h) were included for comparison. **g**, Survival of wild-type (*n* = 53 total; 22 females, 31 males) and *Xrcc4* nKO (*n* = 46 total; 21 females, 25 males) mice. *P*-value was determined by the *Log*-rank Mantel-Cox test. **h**, Locomotor activity (distance traveled in arbitrary units (A.U.), *left*) and quantification of time spent near the walls (*right*) for young (4–6 months) and middle-aged (10–12 months) wild-type or *Xrcc4* nKO mice (*n* = 9–13 mice per genotype), respectively. Data represent mean ± s.d.; one-way ANOVA and Tukey’s *post-hoc* multiple comparison test. **i**, Quantification of errors made during the radial arm water maze training and testing phases by middle-aged (10–12 months) wild-type or *Xrcc4* nKO mice (*n* = 9–11 mice per genotype). Data represent mean ± s.d; two-way ANOVA with Šídák’s *post-hoc* test.

Accumulation of DNA damage has been implicated in the etiology of neuroinflammation and neurodegenerative diseases^1,12,14,25^. By bulk RNA-seq, however, the cortical transcriptome of middle-aged *Xrcc4* nKO showed very few gene expression changes compared to middle-aged wild-type controls. Overall, only 138 genes were differentially expressed (fold-change ≥1.5, adjusted *P*-value < 0.05; **Supplementary Table 1**) and we did not observe altered expression of genes associated with neuroinflammation, including cytokines, inflammatory signaling pathways, or nucleic acid sensors such as *Cgas* or *Tlr9* (**Extended Data Fig. 4a**)^14,25^. Consistent with the absence of neuroinflammation, *Xrcc4* nKO hippocampus and cerebral cortex did not show evidence of astrocytosis or microglial activation, either at young or old age (**Extended Data Fig. 4b,c)**. Moreover, neuronal ablation of *Xrcc4* did not cause detectable decreases in neuronal number or changes in cortical or hippocampal organization at either young or old age (Fig. 4c–e). Similarly, neuronal C-NHEJ inactivation did not induce the expression of factors associated with apoptosis (active caspase-3), inflammation (pSer727-Stat1, total Stat1), or affect the expression of cell cycle- and senescence-associated proteins (pSer15-p53, p21, p16; lamin A/C or B1, respectively) **(Fig. 4f**). Consistent with the absence of signs of neurodegeneration, lifespan of *Xrcc4* nKO mice was unaffected (**Fig. 4g** and **Extended Data Fig. 4d**). Together, these results demonstrate that endogenous neuronal DSBs are not sufficient to cause neurodegeneration or neuroinflammation, even in the absence of C-NHEJ, which we show is the major neuronal DSB repair pathway (**Fig. 2**).

The formation of transcription-associated neuronal DSBs has been implicated in mouse behavior, memory, and learning^9,10^. Open-field testing, however, did not reveal differences in total distance traveled (**Fig. 4h**, *left*) or time spent in the central open area (**Fig. 4h**, *right*) between either young or middle-aged wild-type or *Xrcc4* nKO mice, indicating normal motor function and anxiety levels in mice with neuronal C-NHEJ inactivation. To determine the potential impact of neuronal DSB accumulation on hippocampus-dependent memory and learning functions, we performed radial arm water maze (RAWM) assays (**Fig. 4i**). RAWM testing did not reveal significant differences in spatial memory function between wild-type and *Xrcc4* nKO mice (**Fig. 4i**). These findings demonstrate that enhanced levels of endogenous DSBs are not sufficient to impair memory or learning function or drive aging-associated cognitive decline.

Our findings show that neuronal DSBs occur spontaneously in open chromatin and active promoters and reveal that neuronal DSB-containing genomic regions cluster intra-chromosomally and undergo loop extrusion. Notably, the results from our study provide several essential and surprising insights that affect commonly held notions of the potential role of neuronal DSBs in brain aging: First, wild-type neurons in the aging mouse brain do not show DSB foci accumulation. This indicates that neurons have evolved sufficient DSB repair capacities that generally prevent the accumulation of DSB foci across their long lifespan. Second, in the absence of C-NHEJ, both the fraction of neurons with DSB foci and the number of DSB foci per neuron increase with age, demonstrating that C-NHEJ mediates most neuronal DSB repair. These latter findings also suggest that, unlike somatic cell types such as B cells^6^, neurons do not have robust alternative end-joining repair mechanisms. Third, we show that the DSB burden caused by endogenous processes in neurons is insufficient to cause brain aging or neurodegeneration, even when C-NHEJ, the primary neuronal DSB repair pathway, is inactivated. The latter findings reveal a unique response of mature postmitotic neurons to DSBs. In contrast, inactivation of C-NHEJ *via Xrcc4* ablation in the germline causes late embryonic lethality associated with extensive apoptosis of early postmitotic neurons^18^. Neurodevelopmental apoptosis may serve as a quality control step to eliminate early postmitotic neurons with DSBs before they can be incorporated into the neuronal network. However, eliminating mature postmitotic neurons with DSBs is likely undesirable, given the limited potential for neurogenesis in the mature brain. Thus, the ‘cost’ associated with mature postmitotic neurons carrying persistent DSBs that arise from endogenous sources is likely more tolerable than their complete loss.

## METHODS

### Mice

All studies were approved by the Institutional Animal Care and Use Committee and the Institutional Biosafety Committee at the University of California, San Francisco. Mice were housed on a 12-h light/dark cycle in a temperature- and humidity-controlled environment, with water and food *ad libitum.* The reporting in this manuscript follows the ARRIVE guidelines^26^. Congenic C57BL/6 mice carrying a floxed *Xrcc4* knockout allele were generated by backcrossing mice with a well-characterized conditional *Xrcc4* knockout allele^16^ (*X4c*; a gift from F. W. Alt, Howard Hughes Medical Institute, Boston Children’s Hospital, Harvard Medical School) to C57BL/6 mice for 11 generations. *X4c* mice on a congenic C57BL/6 background were then crossed to a *calcium/calmodulin-dependent protein kinase II alpha* (*Camk2a*) promoter-driven Cre line (RRID:IMSR_JAX:005359)^17^ on a C57BL/6 background. Both female and male mice were used.

### Cells

SV40 large-T-antigen-transformed female C57BL/6 murine tail fibroblasts were grown in DMEM (Gibco, 11965) containing 8.6% (v/v) fetal bovine serum, 10 mM HEPES (Gibco, 15630), 1 mM sodium pyruvate (Gibco, 11360), 2 mM L-glutamine (Gibco, 25030), 100 U/mL penicillin/streptomycin (Gibco, 15140-122), 0.1 mM MEM non-essential amino acids (Gibco, 11140), and 0.1 mM β-Mercaptoethanol (Sigma, M-3148) on gelatinized dishes at 37 °C, 5% CO_2_ in a standard tissue culture incubator.

### Isolation of nuclei

For fluorescent-activated cell sorting (FACS), nuclei were isolated from pooled cortical and hippocampal tissues as previously described, omitting the fixation step^27^. For ATAC-seq and sBLISS, cortical nuclei were isolated as described^28^. For CUT&RUN, cortical nuclei isolation was performed following the Cell Signaling Technology (CST) kit protocol (CST, 86652). For Hi-C, isolation of nuclei was performed as follows: cortices were incubated in Lysis Buffer 1 (1 mM EDTA, 10% (v/v) glycerol, 0.5% (v/v) IGEPAL CA-630, 0.25% (v/v) Triton X-100, 50 mM HEPES-KOH, pH 7.5, 1× cOmplete™ EDTA-free protease inhibitor cocktail (Sigma-Aldrich, 11873580001)) for 10 min on ice. Homogenization was performed with 20 strokes of a loose pestle and 20 strokes of a tight pestle. Nuclei were fixed with 1.68% (v/v) paraformaldehyde for 10 min at room temperature and quenched with 110 mM glycine for 5 min at room temperature. Nuclei were centrifuged for 10 min at 4 °C at 1,000 × g, washed with ice-cold PBS, and sedimented at 1,000 × g for 10 min at 4 °C. For FACS, isolated nuclei were blocked on ice for 30 min in 3% (v/v) goat serum, 5% (w/v) BSA in PBST (0.1% (w/v) Tween 20 in 1×PBS). Anti-NeuN-Alexa555 antibody was added for 30 min on ice. Nuclei were washed with 5% (w/v) BSA/PBST and centrifuged for 10 min at 3,200 × g at 4 °C. Nuclei were resuspended in 5% (w/v) BSA/PBST containing 1 µg µL^−1^ DAPI (Thermo Fisher Scientific, 62248). DAPI/NeuN-positive neuronal nuclei were sorted using a SONY SH800 flow cytometer.

### RNA-seq

Total RNA was isolated from the cerebral cortex with Trizol (Thermo Fisher Scientific, 15596026), followed by *1*-bromo-*3*-chloropropane extraction, isopropanol precipitation, and washing in 75 % (v/v) ethanol. 1 µg of total RNA per sample was used for RNA-seq library preparation using the Illumina TruSeq stranded mRNA kit (Illumina, 20020595). Paired-end sequencing was performed on a NovaSeq (Illumina). We used the *nf-core* framework best-practice RNA-seq analysis pipeline (*nf-core/rnaseq v.3.6*)^29^ with DESeq2^30^ for sequence processing and RNA-seq data analysis. STAR-fusion^31^ (v.1.10.0) was used to detect gene fusion transcripts (*STAR-Fusion --FusionInspector validate --examine_coding_effect -- denovo_reconstruct*).

### sBLISS

Preparation of sBLISS libraries and sequencing was performed as described^32^. Briefly, 1 × 10^6^ nuclei were isolated from fresh cortex and hippocampus from wild-type or *Xrcc4* nKO mice. Nuclei were fixed with 2% (v/v) formaldehyde in PBS for 10 min and quenched with 125 mM glycine for 5 min at room temperature. After nuclear lysis in Lysis Buffer 2 (150 mM NaCl, 0.3% (w/v) SDS, 10 mM Tris-HCl, pH 8.0) for 1 h at 37 °C and washes in CSTX buffer (1× CutSmart buffer (New England Biolabs (NEB) B60004), 0.1% (v/v) Triton X-100), DSB ends were blunted with the Quick Blunting kit (NEB, E1201L) for 1 h at room temperature. Blunted DSB ends were ligated to sBLISS adaptors for 16–18 h at 16 °C using the T4 DNA ligase kit (Thermo Fisher Scientific, EL0012). Following washes with CSTX buffer, samples were incubated with proteinase K (NEB, P8107S) at 55 °C for 18–24 h. DNA was purified by phenol-chloroform extraction and precipitated overnight at −80 °C in 100% ethanol, 3 M NaOAc, and 20 mg mL^−1^ glycogen. After centrifugation, the DNA pellet was washed with 70% (v/v) ethanol and resuspended in TE buffer. DNA was fragmented using dsDNA fragmentase (NEB, M0348) for 40–50 min at 37 °C to generate 300–800-bp fragments. Fragmented DNA was purified using SPRI beads (103778-488, Bulldog Bio). 100 ng of DNA was used to perform *in vitro* transcription for 15 h at 37 °C with the Megascript T7 transcription kit (Thermo Fisher Scientific, AMB13345). 1 U DNase I (Thermo Fisher Scientific, AM2222) was added to each sample to degrade DNA for 15 min at 37 °C. RNA was purified with RNAclean XP beads (Beckman, A63987). RA3 adaptors were ligated to RNA with the T4 RNA ligase kit (NEB, M0242L) for 2 h at 25 °C. RTP primers were used to initiate reverse transcription with the Superscript IV reverse transcriptase kit (Thermo Fisher Scientific, 18090200) following these conditions: 50 °C for 50 min and 80 °C for 10 min. Libraries were indexed and amplified with RPIX-specific primers and RP1 common primer using the following settings: 98 °C, 30 s; 11 cycles [98 °C, 10 s; 60 °C, 30 s; 65 °C, 45 s], 65 °C, 10 min. Bead purification was performed using SPRI beads (Bulldog Bio, 103778-488). Agilent TapeStation 4200 analysis was used to check libraries prior to sequencing (76 cycles in single-end mode; Illumina Nextseq500). Analysis of sBLISS data was performed by using a published pipeline^32^. Single-base pair DSB sites identified by the sBLISS pipeline were extended by adding 37-bp on each side of the DSB to generate 75-bp fragments. DSB-enriched regions were identified by using *MACS2*^33^ (*macs2 callpeak --nomodel --pvalue 1e-5*). BED files were generated from *MACS2 narrowPeak* output and regions overlapping with known ENCODE blacklisted *mm10* regions were removed^34^. sBLISS consensus peaks were defined as sBLISS peaks present in two of three biological replicates of each genotype in young mice or three of four biological replicates of each genotype in middle-aged mice by using *MSPC*^35^. The genomic distribution of sBLISS *MACS2* peaks was determined by using the *HOMER*^36^ (v4.11.1) *annotatePeaks.pl* function. The proximity of sBLISS peaks to genes was determined by T-Gene^37^ (v.5.5.3). *De novo* motif analysis was performed by using *HOMER* (v.4.11.1). Circos plots were generated with *circlize*^38^ (v.0.4.1.5) in R^39^.

### ATAC-seq

ATAC-seq library preparation and sequencing were performed as described^40^, with slight modifications. In brief, 1 × 10^5^ neuronal nuclei from cortex and hippocampus per sample were isolated and resuspended in ATAC-seq lysis buffer (10 mM NaCl, 3 mM MgCl_2_, 0.1 % (w/v) IGEPAL CA-630, 10 mM Tris-HCl, pH 7.4). Transposition was performed in a 50-µL reaction containing 4.7 µL TDE1 Tagment DNA enzyme (Illumina, 20034198), and the reaction was cleaned up (DNA Clean and Concentrator kit; Zymo, D4034). NEBNext Ultra II master mix (NEB, M0544L), SYBR Green I (Thermo Fisher Scientific, S7563), and 0.5 µM of sample-specific P5/P7 barcoded primer mix (NEB, E7335S, E7730S, and E7600S) were used to add barcodes and amplify the ATAC-seq samples by quantitative and limited PCR. SPRI beads (Bulldog Bio, 103778-488) were used to purify libraries. Library quality and quantity were checked using a High Sensitivity DNA kit (Agilent, 5067-4626) and the Qubit dsDNA HS assay (Fisher Scientific, Q32854). Libraries were sequenced for 38 cycles in paired-end mode on a Nextseq500 (Illumina). We used the *nf-core* framework best-practice ATAC-seq analysis pipeline (*nf-core/atacseq* v.1.2.1) for sequence processing and data analysis^41,42^. In brief, *TrimGalore*^43^ (v.0.6.4_dev) was used to trim adapters, and *BWA*^44^ (v0.7.17) was used to map adapter-trimmed reads to the *mm10* reference assembly. *Picard*^45^ (v2.23.1) was used to mark and remove duplicates. Peak calling was performed by *MACS2*^33^ (v2.2.7.1).

### CUT&RUN

Preparation of CUT&RUN libraries was done with a CUT&RUN assay kit (CST, 86652). Briefly, neuronal nuclei from cortex and hippocampus were fixed with 0.1% (w/v) formaldehyde in PBS for 2 min and the fixation reaction was quenched with 100 mM glycine for 5 min at room temperature. Nuclei were washed in 1× wash buffer containing spermidine and protease inhibitor cocktail, captured with activated Concanavalin A beads, and incubated in antibody binding buffer containing spermidine, protease inhibitor cocktail, and digitonin for 16–18 h at 4 °C. Per CUT&RUN reaction, 1 × 10^5^ nuclei were used with either 0.2 µg of H3K27me3 antibody, 0.2 µg of H3K4me3 antibody, or 1 µg of H3K27ac antibody. To bind antibodies to pAG-MNase, nuclei were incubated in 1× wash buffer containing spermidine, protease inhibitor cocktail, digitonin and pAG-MNase for 1 h at 4 °C. pAG-MNase was activated by adding calcium chloride. After a 30-min incubation at 4 °C, the digestion was terminated with stop buffer (digitonin solution, RNase A) containing 50 pg *S. cerevisiae* spike-in DNA for normalization and incubation for 10 min at 37 °C. Crosslinks were reversed by adding SDS and proteinase K for 2 h at 65 °C. The DNA Clean and Concentrator kit (D4034, Zymo) was used to clean DNA samples. Library preparation was performed with the NEBNext Ultra II DNA Library Prep Kit for Illumina (NEB, E7645L). SPRI beads (Bulldog Bio, 103778-488) were used to clean and size-select the libraries. Quantification and quality checks of libraries were performed by Qubit and Tapestation 4200 analysis. Libraries were sequenced for 38 cycles in paired-end mode on a NextSeq500 (Illumina). The *nf-core* framework CUT&RUN analysis pipeline (v3.0; https://github.com/nf-core/cutandrun/tree/3.0)^41,46^ was used for data analysis. Briefly, adapters were trimmed (*TrimGalore*, v.0.6.6) and adapter-trimmed reads were mapped to the mouse (*mm10*) and spike-in (*S. cerevisiae*, *sacCer3*) genome reference assemblies by *Bowtie2*^47^ (v2.4.4). Duplicates were marked and removed by *Picard* (v2.27.4). Reads were normalized against spike-in. Peak calling was performed by *SEACR*^48^ (v1.3) using stringent settings. Consensus peaks were identified as peaks that were present in all replicates per condition. Genomic distribution of CUT&RUN consensus *SEACR* peaks was determined by using the *HOMER* (v4.11.1) *annotatePeaks.pl* function.

### BigWig visualization

For ATAC-seq and CUT&RUN, bigwig files from each sample per genotype were averaged using *DeepTools*^49^ (v.3.5.3) *bigwigAverage*. Matrices for bigwig plots were generated using *DeepTools computeMatrix reference-point –a 3000 –b 3000 – referencePoint center –missingDataAsZero –smartLabels.* Matrices were plotted by using *DeepTools plotProfile*.

### Quantification of ATAC-seq and CUT&RUN signal

Read counts over peak sets were generated using *DeepTools* (v.3.5.3) *multiBigwigSummary*. Outliers were removed using a ROUT’s test at 1% confidence and statistical tests were performed with GraphPad Prism (v.10.2.3).

### Hi-C

Preparation of Hi-C libraries and sequencing was performed as described^50^. Briefly, 1.5 × 10^6^ nuclei were isolated from the cerebral cortex, washed once in PBS, cross-linked with 1.68% paraformaldehyde in PBS, and incubated for 10 min at room temperature. Fixation was quenched with 116 mM glycine for 5 min at room temperature and 15 min on ice. Nuclei were sedimented at 1,000 × g for 10 min at 4 °C. Crosslinked nuclei were resuspended and incubated for 10 min in ice-cold lysis buffer (10 mM NaCl, 0.2% (v/v) IGEPAL CA-630, 10 mM Tris-HCl, pH 8.0). Lysed nuclei were resuspended and incubated for 30 min at 37 °C in 0.5× DNase I digestion buffer containing 0.5 mM MnCl_2_ and 0.2% (v/v) SDS, followed by incubation for 10 min at 37 °C in 0.5× DNase I digestion buffer containing 0.5 mM MnCl_2_, 2% (v/v) Triton X-100, and 80 μg mL^−1^ RNase A (Thermo Fisher Scientific, EN0531). Chromatin digestion was performed by adding 1.5 U of DNase I (Thermo Fisher Scientific, EN0525) per sample and incubation for 4 min at room temperature. To stop the digestion, 6× stop solution (125 mM EDTA, 2.5% (v/v) SDS) was added. Nuclei were bound by AMPure XP beads (Beckman Coulter, A63880) and end repair was performed by incubating chromatin-bound beads for 1 h at room temperature in 1× T4 ligase buffer with ATP (NEB, B0202), 0.25 mM dNTPs (Thermo Fisher Scientific, FERR0181), 0.045 U T4 DNA polymerase (Thermo Fisher Scientific, P0062) and 0.15 U of Klenow fragment (Thermo Fisher Scientific, EP0052). To stop the reaction, SDS was added to a final concentration of 0.25% (v/v). dA-tailing was performed by incubating the chromatin-bead mixture for 1 hour at 37 °C with 1× NEB Buffer 2 (NEB, B7002), 0.5 mM dATP, 1% (v/v) Triton X-100, and 0.375 U of Klenow fragment (exo–, Thermo Fisher Scientific, EP0422). The dA-tailing reaction was stopped by adding SDS to a final concentration of 0.25% (v/v). Biotinylated bridge adaptors (5’: /5Phos/GCTGAGGGA/iBiodT/C; Bridge adaptor 3′T: CCTCAGCT) were ligated overnight to end-repaired and dA-tailed chromatin at 16 °C (8 μM annealed bridge adaptor with biotin, 8 μM annealed blunt adaptor without biotin, 1×T4 ligase buffer with ATP (NEB, B0202), 5% (w/v) PEG-4000, 0.5% (w/v) Triton X-100, 0.25 U T4 DNA ligase (Thermo Fisher Scientific, EL0012)). To stop the reaction, SDS was added to a final concentration of 0.5% (w/v). Bead-bound nuclei were resuspended in nuclease-free water and AMPure buffer (20% (w/v) PEG-8000, 2.5 M NaCl) and washed twice with 80% (v/v) ethanol. Proximity ligation was performed by *in-situ* phosphorylation of adaptors for 1 h at 37 °C (1× T4 DNA ligase buffer with ATP, 1 U PNK (Thermo Fisher Scientific, EK0032)) and *in-situ* ligation for 4 h at room temperature (1× T4 DNA ligase buffer, 0.03 U T4 DNA ligase). Cross-linking reversal was performed for 16–18 h at 60 °C by incubation in 1× NEB Buffer 2, 1% (v/v) SDS with 1.67 mg mL^−1^ proteinase K. DNA precipitation was performed by incubation with GlycoBlue (Fisher Scientific, AM9516), 0.25 M sodium acetate, and isopropanol overnight at −80 °C. After centrifugation, DNA was resuspended in nuclease-free water and purified using AMPure beads and an 80% (v/v) ethanol wash. Purified DNA was sonicated to 150–500 bp using a Covaris M220 (five cycles; duty cycle: 15%; peak incident power: 450; cycles per burst: 200; set mode: frequency sweeping; processing time: 80 s). Biotin-labeled fragments were captured with streptavidin MyOne C1 beads (Life Technologies, 65001). End-repair was performed using the Fast DNA End Repair Kit (Thermo Fisher Scientific, K0771). Samples were dA-tailed using 1× NEBuffer 2, 1 mM dATP and 0.42 U Klenow fragment (exo–, Thermo Fisher Scientific, EP0422) for 30 min at 37 °C. Ligation of sequencing adaptors was performed by incubating end-repaired DNA with 1× Rapid Ligation Buffer (Thermo Fisher Scientific, K1422) supplemented with 0.4 U T4 DNA ligase, and Illumina Y adaptors (SeqAdapt forward: ACACTCTTTCCCTACACGACGCTCTTCCGATC*T; SeqAdapt reverse:/5Phos/GATCGGAAGAGCACACGTCTGAACTCCAGTCAC) for 30 min at room temperature. The reaction was stopped by adding 0.5 M EDTA. 2× HotStart Ready Mix (Roche, KK2601) was used for PCR amplification. Library clean-up and size selection was done by using AMPure XP beads. Libraries were quantified and analyzed by Qubit and Tapestation 4200 analysis and sequenced for 100 cycles in paired-end mode on a Novaseq (Illumina). Reads were aligned to the *mm10* genome with *bwa-mem2*^51^. Ligation junctions were identified, sorted, and indexed using *pairtools*^52^. PCR and optical duplicates were removed by *pairtools dedup*. Hi-C pairs (UU, RU and UR) were selected with *pairtools select*. Hi-C count matrices were generated using *cooler*^53^ and *juicer*^54^ at multiple resolutions. Differential Hi-C heatmaps were generated using *Juicebox*^55^. Hi-C contact matrices to generate Aggregate Peak Analysis (APA) plots were calculated using *Coolpuppy*^56^ at either 10-kb or 50-kb resolution before being processed with a custom R script for visualization. The fold change of APA plots was calculated between the 3×3 central pixels *versus* the 3×3 pixels of the bottom left corner. To identify loop extrusion, aggregate Hi-C heatmaps were generated using *Coolpuppy* at 10-kb or 50-kb resolution with the *–local* flag.

### Targeted amplicon sequencing

Amplicon-targeted libraries were generated as previously described^24^. Per mouse, 30 ng of genomic DNA from FACS-sorted neuronal nuclei were amplified with Phusion Hot Start II high-fidelity DNA polymerase (Thermo Fisher Scientific, F-549L) to add UMI adapters (40 nM) with the following PCR conditions: 98 °C, 30 s; 4 cycles of 98 °C, 10 s; 62 °C, 6 min; 72 °C, 30 s; 65 °C, 15 min; 95 °C, 15 min. During the 15-min step at 65 °C, the PCR reaction was terminated by the addition of *Streptomyces griseus* protease Type XIV (Sigma-Aldrich, P5147) to a final concentration of 0.06 µg µL^−1^. 10 µL of DNA with UMI adapters were amplified by PCR [98 °C, 3 min; 2–35 cycles of 98 °C for 10 s; 80 °C for 1 s; 72 °C for 30 s; 76 °C for 30 s, with ramping at 0.2 °C s^−1^) Duplicate reactions were performed for each mouse to increase the diversity of UMI sampling in the final libraries. The library clean-up and selection of 275–650-bp amplicons was done using SPRI beads. Computational analysis was conducted as described^24^ using *Debarcer* (v.2.1.4; https://github.com/oicr-gsi/debarcer).

### Immunofluorescence microscopy and quantification

Mice were anesthetized with 200 mg ketamine, 40 mg xylazine kg^−1^, and transcardially perfused with 50 mL PBS. Whole brains were collected, fixed in 4% paraformaldehyde overnight at 4°C, and transferred to a 30 % (w/v) sucrose solution for cryoprotection. Brains were incubated overnight with end-over-end rotation at 4 °C, embedded in OCT compound, and frozen on dry ice. 30-µm thick, free-floating coronal sections were collected using a Leica Cryostat HM550. Free-floating sections were permeabilized in TBS containing 1% (v/v) Triton X-100 for 20 min at 4 °C and blocked for 45 min at room temperature in blocking solution (5% (v/v) normal goat serum, 0.1% (v/v) Tween 20, TBS). Sections were incubated overnight at 4 °C in blocking solution containing primary antibodies. After three washes in TBS containing 0.1% (v/v) Triton X-100, sections were incubated for 1 h at room temperature in blocking solution containing secondary antibodies. After two washes in TBS containing 0.1% (v/v) Triton X-100, sections were stained with DAPI (1 μg mL^−1^ in PBS) for 10 min at room temperature. Sections were washed once and mounted onto glass slides. Sections were imaged using a Nikon spinning disk confocal microscope. DSB response foci were quantified using an automated CellProfiler^57^ pipeline. For each region, two fields of view from one coronal section were imaged, scored, and pooled to get mean values for each mouse. Cortical width, DAPI^+^, NeuN^+^, and GFAP^+^ area was measured using the *ImageJ* Analyze tool. Two coronal sections were averaged per mouse.

### Immunoblotting

The cortex and hippocampus were collected, snap-frozen, reduced to powder in liquid nitrogen, and homogenized in ice-cold RIPA buffer (150 mM NaCl, 1% (v/v) NP-40, 0.5% (w/v) sodium deoxycholate, 0.1% (w/v) SDS, 50 mM Tris-HCl, pH 8.0, 1× protease/phosphatase inhibitor cocktail). 2 mM MgCl_2_ and 479 U mL^−1^ Benzonase (Sigma-Aldrich, E1014) were added, and samples were incubated for 1 h at 4 °C. Samples were centrifuged for 10 min at 10,000 × g to remove insoluble material and the supernatant was recovered. Protein concentration was determined using a BCA Protein Assay (Pierce). Protein extracts were diluted in sample buffer (100 mM DTT, 2% (w/v) SDS, 10% (w/v) glycerol, 0.002% (w/v) bromophenol blue, 50 mM Tris-HCl, pH 6.8), denatured for 5 min at 95 °C, separated on NuPAGE 4–12% Bis-Tris protein gels (Life Technologies), and transferred to PVDF membranes (Millipore). Membranes were blocked with 3% (w/v) BSA in TBST (0.1% (v/v) Tween 20 in TBS) for 30 min at room temperature and incubated with primary antibodies overnight at 4 °C. Following three TBST washes, membranes were incubated for 1 h at room temperature with horseradish-peroxidase-conjugated secondary antibodies. Membranes were washed, incubated with ECL substrate, and imaged (ChemiDoc Imaging System, Biorad).

### Lifespan analysis

Animals that were euthanized for tissue collection were censored from the lifespan analysis. The *Log*-rank Mantel-Cox test was used for statistical analysis.

### Open field test

Mice were placed into the center of a square (40 cm × 40 cm) open chamber (Kinder Scientific) and allowed to explore freely for 10 min. The open chamber did not contain any cues or stimuli. Motion metrics were analyzed using infrared photobeam breaks and *MotorMonitor* software (Kinder Scientific).

### Radial arm water maze

The radial arm water maze (RAWM) test was performed as described^58,59^. The maze consists of a circular swimming pool with six arms. The goal arm location with the platform remained constant while the start arm was changed at each trial. Spatial cues were placed on the room walls. Entry into an arm that was not the goal arm was scored as an error, and errors were averaged per training block (three consecutive trials). On day one (training phase), mice were trained for 12 trials to find the goal arm, with trials alternating between a visible and hidden platform (blocks 1–4). After a one-hour break, learning was assessed for three trials (block 5) using a hidden platform. On day two (testing phase), mice were tested for 15 trials (blocks 6–10) using a hidden platform. Investigators were blinded to genotype when scoring.

### Statistical analysis

No statistical methods were used to predetermine the sample size. Samples for image acquisition and analysis, and mice for behavioral testing were randomized, and investigators were blinded to allocation during the experiment and outcome assessment. Details on statistical analysis are listed in the figure legends. Briefly, comparisons between two groups were performed by unpaired, two-tailed *t*-test, and comparisons across multiple groups were done by using one-way ANOVA with Tukey’s *post-hoc* multiple comparisons test. RAWM data was analyzed as previously published^59^, using two-way ANOVA analysis with Šídák’s *post-hoc* test. RAWM data were found to be normally distributed by the Shapiro-Wilk test. Graphs were generated using GraphPad Prism (v.10.2.3).

## ACKNOWLEDGEMENTS

We thank Schwer lab and Mullins lab members for helpful discussions. We thank Sam Lord for advice and assistance with imaging experiments. We thank Coline Arnould and Nadav Ahituv for help with sBLISS protocols. This work was supported by the UCSF Program for Breakthrough Biomedical Research (which is partially funded by the Sandler Foundation), the Carol & Gene Ludwig Family Foundation, the UCSF Bakar Aging Research Institute, and NIH R01 AG064363 (B.S). S.C. was supported by a Sullivan Postdoctoral Fellowship. R.D.M. is an investigator of the Howard Hughes Medical Institute. Sequencing was performed, in part, at the UCSF CAT, supported by UCSF PBBR, RRP IMIA, and NIH 1S10OD028511-01 grants. This study was further supported, in part, by the HDFCCC Laboratory for Cell Analysis Shared Resource Facility through NIH grant P30CA082103. Fig. 1a and Fig. 4i were created with content licensed from *BioRender*.

## AUTHOR CONTRIBUTIONS

B.S. conceived the study. S.C., L.C., and B.S. designed experiments and planned the study. S.C., L.C., K.J.B.P, A.B., A.C., and B.S. performed research. V.R. provided experimental protocols and reagents. S.C., L.C., K.J.B.P, A.C., and B.S. analyzed data. S.A.V., R.D.M., and B.S. provided intellectual guidance and resources and supervised research. S.C., L.C., and B.S. wrote the manuscript with input from all authors.

## EXTENDED DATA

**Extended Data Figure 1.**
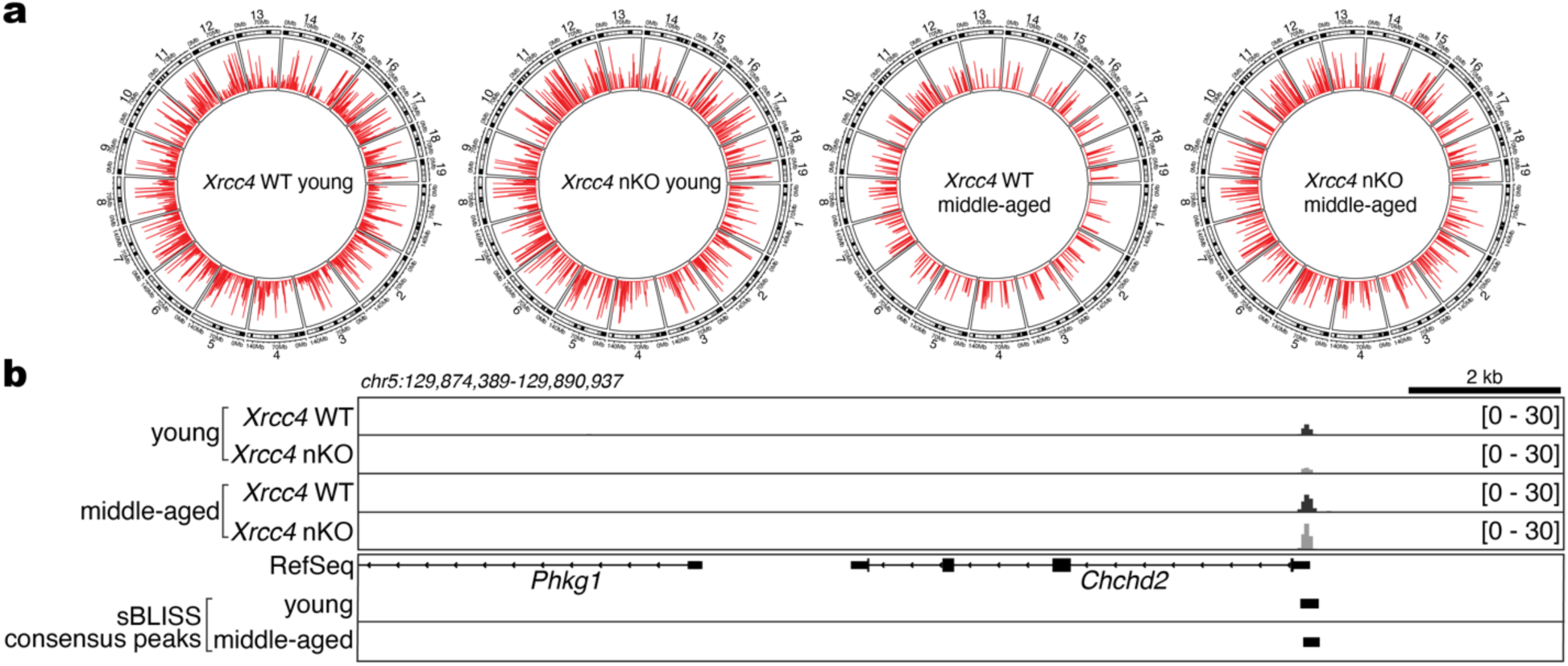
Mapping of endogenous DSBs by sBLISS. **a**, Circos plots of the mouse genome separated into chromosomes showing the genomic distribution of DSB consensus peaks in the cerebral cortex from young (*n* = 3 mice per genotype; 3 months) and middle-aged (*n* = 4 mice per genotype; 13.1 ± 1.1 months) wild-type or *Xrcc4* nKO mice. **b,** Genomic tracks showing aggregate sBLISS signal from the cerebral cortex of young (*n* = 3 mice per genotype; 3 months) and middle-aged (*n* = 4 mice per genotype; 13.1 ± 1.1 months) wild-type or *Xrcc4* nKO mice. Consensus peaks identified in the cerebral cortex of young and old mice are indicated by black bars.

**Extended Data Figure 2.**
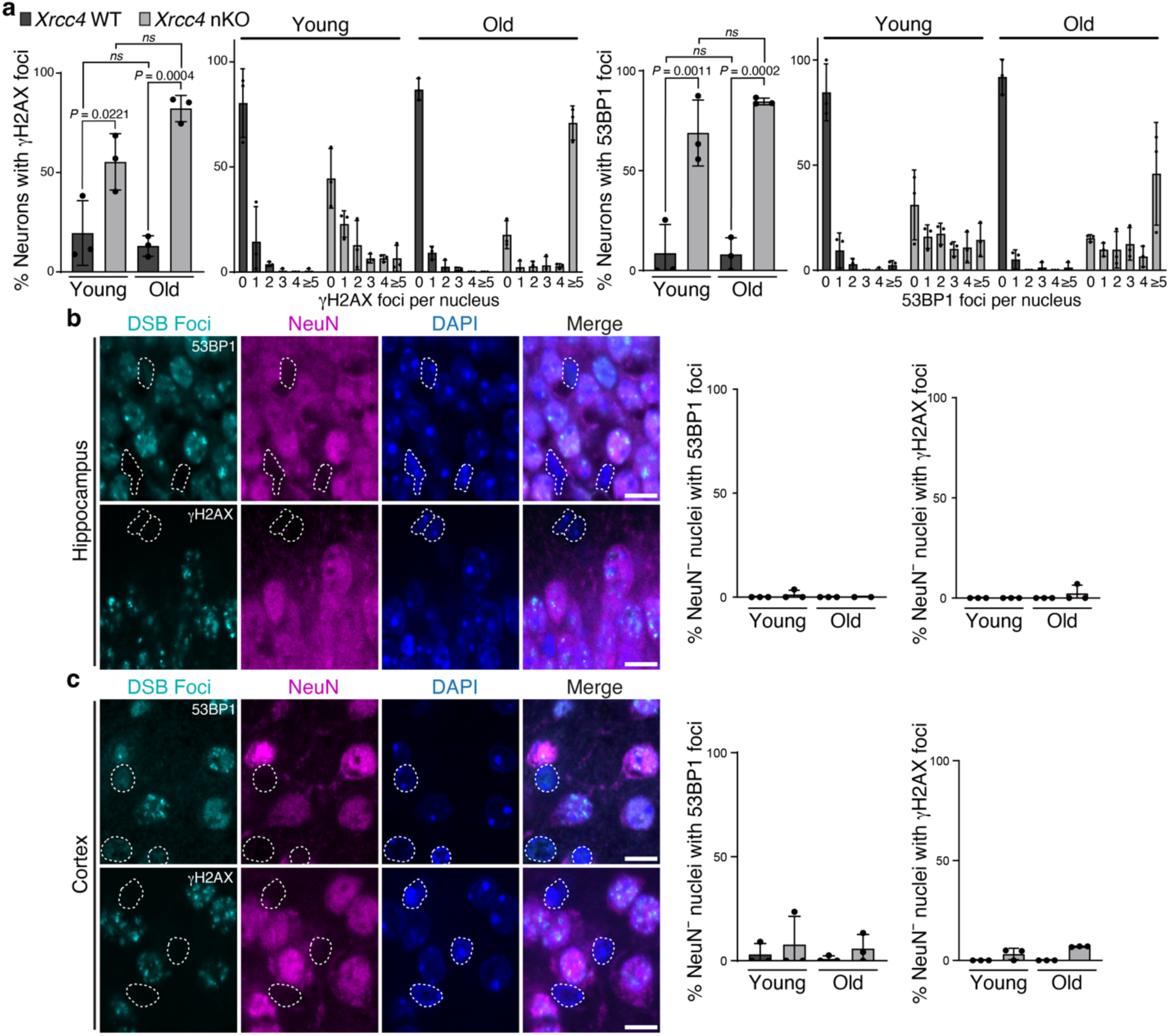
Quantification of DSB response foci. **a**, Quantification of cortical NeuN^+^ neurons with DSB response foci and distribution of the number of foci per neuron based on γH2AX (*left graphs*) or 53BP1 (*right graphs*) staining. Data represent mean ± s.d from *n* = 3 mice per genotype; individual points show the mean from two fields per mouse. *P*-values were determined by one-way ANOVA and Tukey’s *post-hoc* multiple comparisons test; *ns*, not significant. **b**, Representative images of DSB response foci marked by 53BP1 (*top*) and γH2AX (*bottom*) in hippocampal (*Dentate gyrus)* sections of *Xrcc4* nKO mice. Quantifications of non-neuronal cells (DAPI^+^ NeuN^-^) with DSB response foci (53BP1 and γH2AX) are shown on the right. **c**, Representative images of DSB response foci marked by 53BP1 (*top*) and γH2AX (*bottom*) in cortical sections of *Xrcc4* nKO mice. Quantifications of non-neuronal cells (DAPI^+^ NeuN^-^) with DSB response foci (53BP1 and γH2AX) are shown on the right. In **b** and **c**, non-neuronal cells (DAPI^+^ NeuN^-^) are indicated by dotted white lines; scale bars, 10 μm. No significant differences were detected (one-way ANOVA with Tukey’s *post-hoc* multiple comparisons test, *n* = 3 mice per genotype).

**Extended Data Figure 3.**
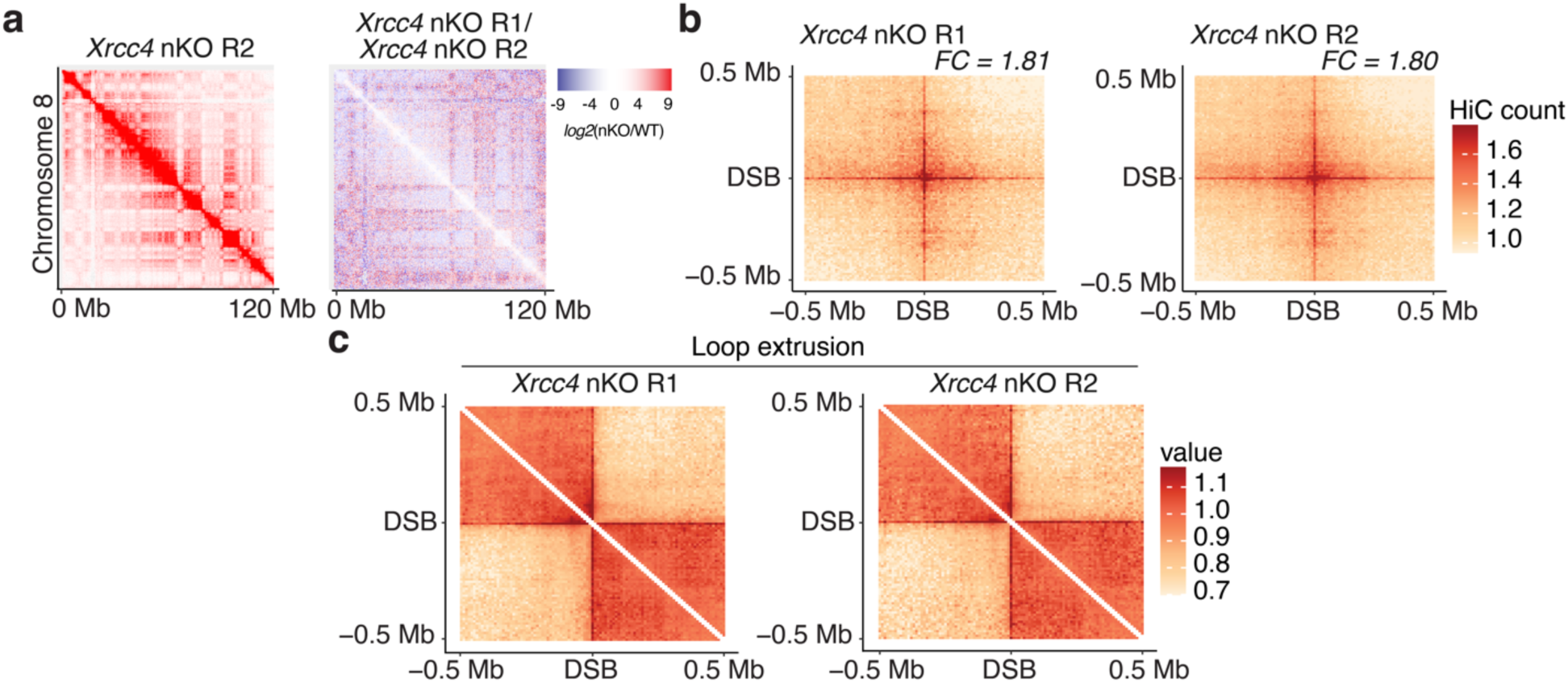
Analysis of 3-D genome organization. **a**, Hi-C contact matrix of chromosome 8 in *Xrcc4* nKO replicate 2 (R2, *left*) and differential Hi-C contact matrix (*log2*[*Xrcc4* nKO R2/WT], *right*). **b**, Mean aggregate peak analysis of intra-chromosomal DSB consensus peak interactions plotted on a 500-kb window (10-kb resolution) in *Xrcc4* nKO cerebral cortex (biological replicates R1 and R2). *FC*, fold change calculated between the central pixel and a square of 3×3 pixels on the bottom left corner of the matrix. **c**, Aggregate Hi-C contact matrix plotted on a 500-kb window (10-kb resolution) centered on sBLISS consensus peaks in *Xrcc4* nKO cerebral cortex (biological replicates *Xrcc4* nKO R1 and *Xrcc4* nKO R2).

**Extended Data Figure 4.**
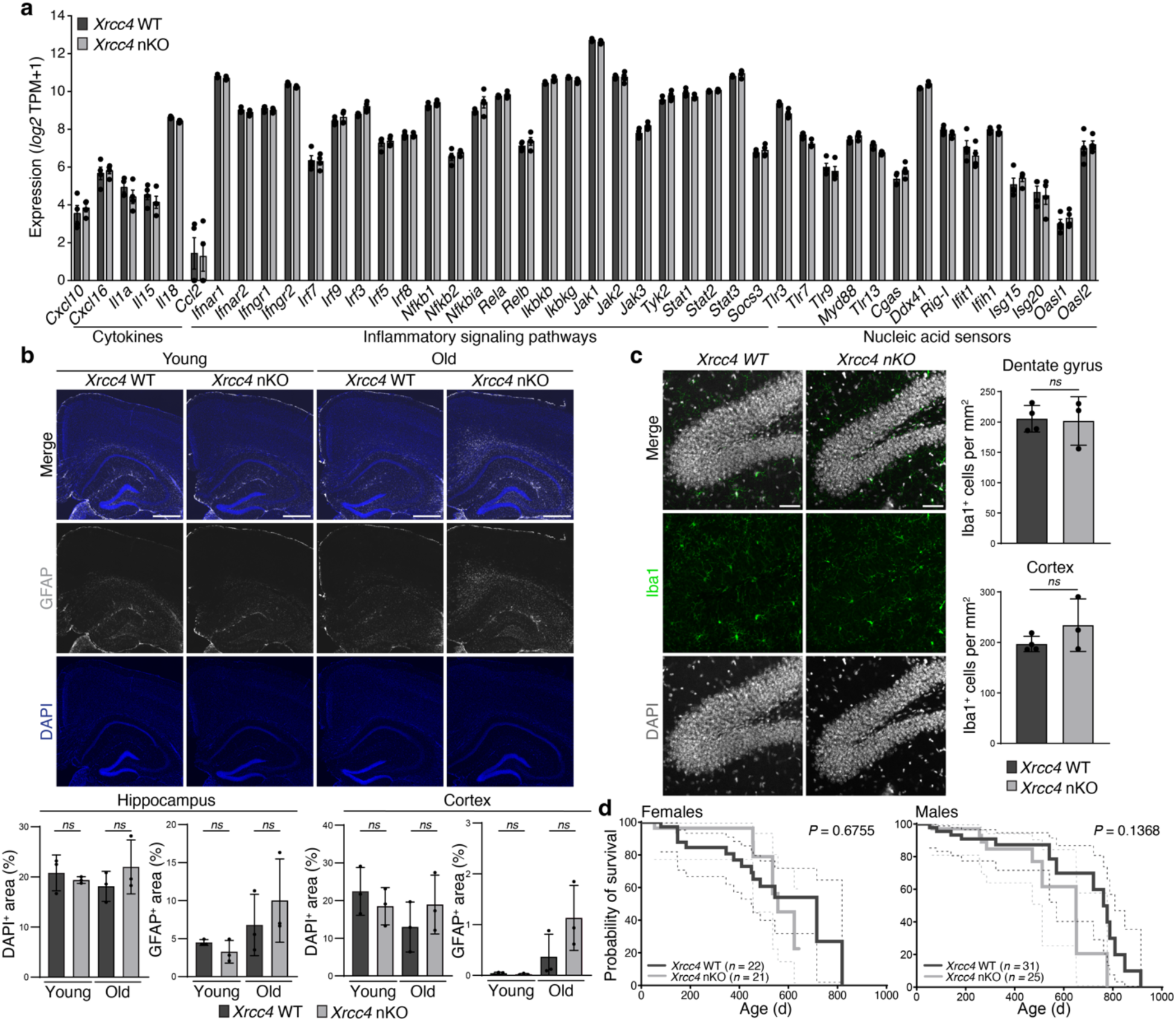
Assessment of neuroinflammatory markers, cortical and hippocampal integrity, and lifespan in mice with neuron-specific C-NHEJ inactivation. **a,** Expression of the indicated neuroinflammation-associated genes in the cerebral cortex of 15.9 ± 1 month-old mice (*n* = 4 per genotype) measured by RNA-seq. Data represent mean ± s.e.m. No significant changes were detected between wild-type and *Xrcc4* nKO (multiple unpaired *t*-tests with two-stage step-up (Benjamini, Krieger, and Yekutieli) multiple comparison and *FDR* = 1%). **b,** Representative images (*top*) of GFAP- and DAPI-stained coronal brain sections (cortex and hippocampus) from young (6.1 ± 1.3 months) and old (21.4 ± 2.5 months) wild-type or *Xrcc4* nKO mice (scale bars, 500 μm). Quantification (*bottom*) of DAPI- or GFAP-positive area in the hippocampus (*Dentate gyrus*, *Cornu ammonis* (CA)1, CA2, and CA3; *left*) and cortex (*right*) of young (6.1 ± 1.3 months) and old (21.4 ± 2.5 months) wild-type or *Xrcc4* nKO mice (*n* = 3 mice per genotype). Data represent mean ± s.d; individual points show the average from two sections per mouse; one-way ANOVA and Tukey’s *post-hoc* multiple comparisons test; *ns*, not significant **c**, Representative images (*left*) and quantification (*right*) of Iba1-positive cells in *Dentate gyrus* sections of young (4 months) wild-type or *Xrcc4* nKO mice (*n* = 3 mice per genotype); scale bars, 50 μm. Unpaired, two-tailed *t*-test. **d,** Survival of wild-type and *Xrcc4* nKO females (*n* = 43, *left*) and males (*n* = 56, *right*). *P-*values were determined by a two-sided *Log*-rank Mantel-Cox test. Dashed lines denote the 95% confidence interval as upper and lower bands for each curve.

## SUPPLEMENTARY MATERIAL

**Supplementary Table 1**. List of upregulated and downregulated transcripts in *Xrcc4* nKO cortical tissue as identified by RNA-seq.

## REFERENCES

1. Wilson, D. M., Cookson, M. R., Bosch, L. V. D., Zetterberg, H., Holtzman, D. M. & Dewachter, I. Hallmarks of neurodegenerative diseases. Cell 186, 693–714 (2023).

2. López-Otín, C., Blasco, M. A., Partridge, L., Serrano, M. & Kroemer, G. Hallmarks of aging: An expanding universe. Cell 186, 243–278 (2023).

3. Szilard, L. On the nature of the aging process. Proceedings of the National Academy of Sciences 45, 30–45 (1959).

4. Miller, M. B., Reed, H. C. & Walsh, C. A. Brain Somatic Mutation in Aging and Alzheimer’s Disease. Annu Rev Genomics Hum Genet 22, 239–256 (2021).

5. Alt, F. W., Zhang, Y., Meng, F.-L., Guo, C. & Schwer, B. Mechanisms of programmed DNA lesions and genomic instability in the immune system. Cell 152, 417–429 (2013).

6. Yan, C. T., Boboila, C., Souza, E. K., Franco, S., Hickernell, T. R., Murphy, M., Gumaste, S., Geyer, M., Zarrin, A. A., Manis, J. P., Rajewsky, K. & Alt, F. W. IgH class switching and translocations use a robust non-classical end-joining pathway. Nature 449, 478–482 (2007).

7. Alt, F. W. & Schwer, B. DNA double-strand breaks as drivers of neural genomic change, function, and disease. DNA Repair (Amst) 71, 158–163 (2018).

8. Delint-Ramirez, I., Konada, L., Heady, L., Rueda, R., Jacome, A. S. V., Marlin, E., Marchioni, C., Segev, A., Kritskiy, O., Yamakawa, S., Reiter, A. H., Tsai, L.-H. & Madabhushi, R. Calcineurin dephosphorylates topoisomerase IIβ and regulates the formation of neuronal-activity-induced DNA breaks. Molecular Cell (2022). doi:10.1016/j.molcel.2022.09.012

9. Suberbielle, E., Sanchez, P. E., Kravitz, A. V., Wang, X., Ho, K., Eilertson, K., Devidze, N., Kreitzer, A. C. & Mucke, L. Physiologic brain activity causes DNA double-strand breaks in neurons, with exacerbation by amyloid-β. Nat Neurosci 16, 613–621 (2013).

10. Madabhushi, R., Gao, F., Pfenning, A. R., Pan, L., Yamakawa, S., Seo, J., Rueda, R., Phan, T. X., Yamakawa, H., Pao, P.-C., Stott, R. T., Gjoneska, E., Nott, A., Cho, S., Kellis, M. & Tsai, L.-H. Activity-Induced DNA Breaks Govern the Expression of Neuronal Early-Response Genes. Cell 161, 1592–1605 (2015).

11. Pollina, E. A., Gilliam, D. T., Landau, A. T., Lin, C., Pajarillo, N., Davis, C. P., Harmin, D. A., Yap, E.-L., Vogel, I. R., Griffith, E. C., Nagy, M. A., Ling, E., Duffy, E. E., Sabatini, B. L., Weitz, C. J. & Greenberg, M. E. A NPAS4-NuA4 complex couples synaptic activity to DNA repair. Nature 614, 732–741 (2023).

12. Yang, J.-H., Hayano, M., Griffin, P. T., Amorim, J. A., Bonkowski, M. S., Apostolides, J. K., Salfati, E. L., Blanchette, M., Munding, E. M., Bhakta, M., Chew, Y. C., Guo, W., Yang, X., Maybury-Lewis, S., Tian, X., Ross, J. M., Coppotelli, G., Meer, M. V., Rogers-Hammond, R., Vera, D. L., Lu, Y. R., Pippin, J. W., Creswell, M. L., Dou, Z., Xu, C., Mitchell, S. J., Das, A., O’Connell, B. L., Thakur, S., Kane, A. E., Su, Q., Mohri, Y., Nishimura, E. K., Schaevitz, L., Garg, N., Balta, A.-M., Rego, M. A., Gregory-Ksander, M., Jakobs, T. C., Zhong, L., Wakimoto, H., El Andari, J., Grimm, D., Mostoslavsky, R., Wagers, A. J., Tsubota, K., Bonasera, S. J., Palmeira, C. M., Seidman, J. G., Seidman, C. E., Wolf, N. S., Kreiling, J. A., Sedivy, J. M., Murphy, G. F., Green, R. E., Garcia, B. A., Berger, S. L., Oberdoerffer, P., Shankland, S. J., Gladyshev, V. N., Ksander, B. R., Pfenning, A. R., Rajman, L. A. & Sinclair, D. A. Loss of epigenetic information as a cause of mammalian aging. Cell 186, 305–326.e27 (2023).

13. Dileep, V., Boix, C. A., Mathys, H., Marco, A., Welch, G. M., Meharena, H. S., Loon, A., Jeloka, R., Peng, Z., Bennett, D. A., Kellis, M. & Tsai, L.-H. Neuronal DNA double-strand breaks lead to genome structural variations and 3D genome disruption in neurodegeneration. Cell 186, 4404–4421.e20 (2023).

14. Welch, G. M., Boix, C. A., Schmauch, E., Davila-Velderrain, J., Victor, M. B., Dileep, V., Bozzelli, P. L., Su, Q., Cheng, J. D., Lee, A., Leary, N. S., Pfenning, A. R., Kellis, M. & Tsai, L.-H. Neurons burdened by DNA double-strand breaks incite microglia activation through antiviral-like signaling in neurodegeneration. Sci Adv 8, eabo4662 (2022).

15. Iyama, T. & Wilson, D. M. DNA repair mechanisms in dividing and non-dividing cells. DNA Repair 12, 620–636 (2013).

16. Yan, C. T., Kaushal, D., Murphy, M., Zhang, Y., Datta, A., Chen, C., Monroe, B., Mostoslavsky, G., Coakley, K., Gao, Y., Mills, K. D., Fazeli, A. P., Tepsuporn, S., Hall, G., Mulligan, R., Fox, E., Bronson, R., De Girolami, U., Lee, C. & Alt, F. W. XRCC4 suppresses medulloblastomas with recurrent translocations in p53-deficient mice. Proceedings of the National Academy of Sciences 103, 7378–7383 (2006).

17. Tsien, J. Z., Chen, D. F., Gerber, D., Tom, C., Mercer, E. H., Anderson, D. J., Mayford, M., Kandel, E. R. & Tonegawa, S. Subregion- and Cell Type–Restricted Gene Knockout in Mouse Brain. Cell 87, 1317–1326 (1996).

18. Gao, Y., Sun, Y., Frank, K. M., Dikkes, P., Fujiwara, Y., Seidl, K. J., Sekiguchi, J. M., Rathbun, G. A., Swat, W., Wang, J., Bronson, R. T., Malynn, B. A., Bryans, M., Zhu, C., Chaudhuri, J., Davidson, L., Ferrini, R., Stamato, T., Orkin, S. H., Greenberg, M. E. & Alt, F. W. A Critical Role for DNA End-Joining Proteins in Both Lymphogenesis and Neurogenesis. Cell 95, 891–902 (1998).

19. Suske, G. NF-Y and SP transcription factors — New insights in a long-standing liaison. Biochimica et Biophysica Acta (BBA) - Gene Regulatory Mechanisms 1860, 590–597 (2017).

20. Brickner, J. R., Garzon, J. L. & Cimprich, K. A. Walking a tightrope: The complex balancing act of R-loops in genome stability. Molecular Cell 82, 2267–2297 (2022).

21. Popp, H. D., Brendel, S., Hofmann, W.-K. & Fabarius, A. Immunofluorescence Microscopy of γH2AX and 53BP1 for Analyzing the Formation and Repair of DNA Double-strand Breaks. JoVE 56617 (2017). doi:10.3791/56617

22. Aymard, F., Aguirrebengoa, M., Guillou, E., Javierre, B. M., Bugler, B., Arnould, C., Rocher, V., Iacovoni, J. S., Biernacka, A., Skrzypczak, M., Ginalski, K., Rowicka, M., Fraser, P. & Legube, G. Genome wide mapping of long range contacts unveils DNA Double Strand Breaks clustering at damaged active genes. Nat Struct Mol Biol 24, 353–361 (2017).

23. Arnould, C., Rocher, V., Finoux, A.-L., Clouaire, T., Li, K., Zhou, F., Caron, P., Mangeot, P. E., Ricci, E. P., Mourad, R., Haber, J. E., Noordermeer, D. & Legube, G. Loop extrusion as a mechanism for formation of DNA damage repair foci. Nature 590, 660–665 (2021).

24. Ståhlberg, A., Krzyzanowski, P. M., Egyud, M., Filges, S., Stein, L. & Godfrey, T. E. Simple multiplexed PCR-based barcoding of DNA for ultrasensitive mutation detection by next-generation sequencing. Nat Protoc 12, 664–682 (2017).

25. Jovasevic, V., Wood, E. M., Cicvaric, A., Zhang, H., Petrovic, Z., Carboncino, A., Parker, K. K., Bassett, T. E., Moltesen, M., Yamawaki, N., Login, H., Kalucka, J., Sananbenesi, F., Zhang, X., Fischer, A. & Radulovic, J. Formation of memory assemblies through the DNA-sensing TLR9 pathway. Nature 1–9 (2024). doi:10.1038/s41586-024-07220-7

26. Kilkenny, C., Browne, W. J., Cuthill, I. C., Emerson, M. & Altman, D. G. Improving Bioscience Research Reporting: The ARRIVE Guidelines for Reporting Animal Research. PLOS Biology 8, e1000412 (2010).

27. Eremenko, E., Golova, A., Stein, D., Einav, M., Khrameeva, E. & Toiber, D. FACS-based isolation of fixed mouse neuronal nuclei for ATAC-seq and Hi-C. STAR Protocols 2, 100643 (2021).

28. Corces, R., Greenleaf, W. J. & Chang, H. Y. Isolation of nuclei from frozen tissue for ATAC-seq and other epigenomic assays. (2019). At <https://www.protocols.io/view/isolation-of-nuclei-from-frozen-tissue-for-atac-se-6t8herw>

29. Patel, H., Ewels, P., Peltzer, A., Hammarén, R., Botvinnik, O., Sturm, G., Moreno, D., Vemuri, P., silviamorins, Pantano, L., Binzer-Panchal, M., Kelly, G., FriederikeHanssen, Garcia, M. U., bot, nf-core, Cheshire, C., rfenouil, Espinosa-Carrasco, J., marchoeppner, Zhou, P., Gabernet, G., Mertes, C., Straub, D., Hörtenhuber, M., Tommaso, P. D., F, S., Hall, G., Panneerselvam, S., OMeally, D. & jun-wan. nf-core/rnaseq: nf-core/rnaseq v3.6 - Platinum Platypus. (2022). doi:10.5281/zenodo.6327553

30. Love, M. I., Huber, W. & Anders, S. Moderated estimation of fold change and dispersion for RNA-seq data with DESeq2. Genome Biol 15, 550 (2014).

31. Haas, B. J., Dobin, A., Li, B., Stransky, N., Pochet, N. & Regev, A. Accuracy assessment of fusion transcript detection via read-mapping and de novo fusion transcript assembly-based methods. Genome Biology 20, 213 (2019).

32. Bouwman, B. A. M., Agostini, F., Garnerone, S., Petrosino, G., Gothe, H. J., Sayols, S., Moor, A. E., Itzkovitz, S., Bienko, M., Roukos, V. & Crosetto, N. Genome-wide detection of DNA double-strand breaks by in-suspension BLISS. Nat Protoc 15, 3894–3941 (2020).

33. Zhang, Y., Liu, T., Meyer, C. A., Eeckhoute, J., Johnson, D. S., Bernstein, B. E., Nusbaum, C., Myers, R. M., Brown, M., Li, W. & Liu, X. S. Model-based analysis of ChIP-Seq (MACS). Genome Biol 9, R137 (2008).

34. Amemiya, H. M., Kundaje, A. & Boyle, A. P. The ENCODE Blacklist: Identification of Problematic Regions of the Genome. Sci Rep 9, 9354 (2019).

35. Jalili, V., Matteucci, M., Masseroli, M. & Morelli, M. J. Using combined evidence from replicates to evaluate ChIP-seq peaks. Bioinformatics 31, 2761–2769 (2015).

36. Heinz, S., Benner, C., Spann, N., Bertolino, E., Lin, Y. C., Laslo, P., Cheng, J. X., Murre, C., Singh, H. & Glass, C. K. Simple combinations of lineage-determining transcription factors prime cis-regulatory elements required for macrophage and B cell identities. Mol Cell 38, 576–589 (2010).

37. O’Connor, T., Grant, C. E., Bodén, M. & Bailey, T. L. T-Gene: improved target gene prediction. Bioinformatics 36, 3902–3904 (2020).

38. Gu, Z., Gu, L., Eils, R., Schlesner, M. & Brors, B. circlize Implements and enhances circular visualization in R. Bioinformatics 30, 2811–2812 (2014).

39. R Core Team. R: A Language and Environment for Statistical Computing. (R Foundation for Statistical Computing, 2021). at <https://www.R-project.org/>

40. Brunton, H., Garner, I. M., Bailey, U.-M., Upstill-Goddard, R. & Bailey, P. J. Using Chromatin Accessibility to Delineate Therapeutic Subtypes in Pancreatic Cancer Patient-Derived Cell Lines. STAR Protoc 1, 100079 (2020).

41. Ewels, P. A., Peltzer, A., Fillinger, S., Patel, H., Alneberg, J., Wilm, A., Garcia, M. U., Di Tommaso, P. & Nahnsen, S. The nf-core framework for community-curated bioinformatics pipelines. Nat Biotechnol 38, 276–278 (2020).

42. Patel, H., Espinosa-Carrasco, J., Langer, B., Ewels, P., bot, nf-core, Garcia, M. U., Syme, R., Peltzer, A., Talbot, A., Behrens, D., Gabernet, G., Jin, M., Hörtenhuber, M., Rodriguez, J. G., Menden, K. & An, Ö. nf-core/atacseq: [2.1.2] - 2022-08-07. (2023). doi:10.5281/zenodo.8222875

43. Babraham Bioinformatics - Trim Galore! At <https://www.bioinformatics.babraham.ac.uk/projects/trim_galore/>

44. Li, H. & Durbin, R. Fast and accurate short read alignment with Burrows-Wheeler transform. Bioinformatics 25, 1754–1760 (2009).

45. Picard Tools - By Broad Institute. at <https://broadinstitute.github.io/picard/>

46. Cheshire, C., charlotte-west, bot, nf-core, Patel, H., Ladd, D., Thiery, A., Fields, C., Deu-Pons, J., Ewels, P., Menden, K. & tamara-hodgetts. nf-core/cutandrun: nf-core/cutandrun v3.1 Lead Rooster. (2023). doi:10.5281/zenodo.7715959

47. Langmead, B. & Salzberg, S. L. Fast gapped-read alignment with Bowtie 2. Nat Methods 9, 357–359 (2012).

48. Meers, M. P., Tenenbaum, D. & Henikoff, S. Peak calling by Sparse Enrichment Analysis for CUT&RUN chromatin profiling. Epigenetics & Chromatin 12, 42 (2019).

49. Ramírez, F., Ryan, D. P., Grüning, B., Bhardwaj, V., Kilpert, F., Richter, A. S., Heyne, S., Dündar, F. & Manke, T. deepTools2: a next generation web server for deep-sequencing data analysis. Nucleic Acids Research 44, W160–W165 (2016).

50. Ramani, V., Cusanovich, D. A., Hause, R. J., Ma, W., Qiu, R., Deng, X., Blau, C. A., Disteche, C. M., Noble, W. S., Shendure, J. & Duan, Z. Mapping three-dimensional genome architecture through in situ DNase Hi-C. Nat Protoc 11, 2104–2121 (2016).

51. Vasimuddin, Md., Misra, S., Li, H. & Aluru, S. Efficient Architecture-Aware Acceleration of BWA-MEM for Multicore Systems. in 2019 IEEE International Parallel and Distributed Processing Symposium (IPDPS) 314–324 (2019). doi:10.1109/IPDPS.2019.00041

52. Open2C, Abdennur, N., Fudenberg, G., Flyamer, I. M., Galitsyna, A. A., Goloborodko, A., Imakaev, M. & Venev, S. V. Pairtools: from sequencing data to chromosome contacts. 2023.02.13.528389 Preprint at 10.1101/2023.02.13.528389 (2023)

53. Abdennur, N. & Mirny, L. A. Cooler: scalable storage for Hi-C data and other genomically labeled arrays. Bioinformatics 36, 311–316 (2020).

54. Durand, N. C., Shamim, M. S., Machol, I., Rao, S. S. P., Huntley, M. H., Lander, E. S. & Aiden, E. L. Juicer Provides a One-Click System for Analyzing Loop-Resolution Hi-C Experiments. Cell Syst 3, 95–98 (2016).

55. Robinson, J. T., Turner, D., Durand, N. C., Thorvaldsdóttir, H., Mesirov, J. P. & Aiden, E. L. Juicebox.js Provides a Cloud-Based Visualization System for Hi-C Data. Cell Syst 6, 256–258.e1 (2018).

56. Flyamer, I. M., Illingworth, R. S. & Bickmore, W. A. Coolpup.py: versatile pile-up analysis of Hi-C data. Bioinformatics 36, 2980–2985 (2020).

57. Stirling, D. R., Swain-Bowden, M. J., Lucas, A. M., Carpenter, A. E., Cimini, B. A. & Goodman, A. CellProfiler 4: improvements in speed, utility and usability. BMC Bioinformatics 22, 433 (2021).

58. Alamed, J., Wilcock, D. M., Diamond, D. M., Gordon, M. N. & Morgan, D. Two-day radial-arm water maze learning and memory task; robust resolution of amyloid-related memory deficits in transgenic mice. Nat Protoc 1, 1671–1679 (2006).

59. Schroer, A. B., Ventura, P. B., Sucharov, J., Misra, R., Chui, M. K. K., Bieri, G., Horowitz, A. M., Smith, L. K., Encabo, K., Tenggara, I., Couthouis, J., Gross, J. D., Chan, J. M., Luke, A. & Villeda, S. A. Platelet factors attenuate inflammation and rescue cognition in ageing. Nature 620, 1071–1079 (2023).

